# Identification of INSR protein structural differences in Diabetes Mellitus type 2 subjects from protein sequences

**DOI:** 10.1101/2021.08.17.456584

**Authors:** Peyton Carroll

## Abstract

In this paper, we constructed several models of the Insulin Receptor protein(abbreviated INSR) from protein sequences from pathogenic and benign variants of type 2 Diabetes Mellitus patients. Through modeling these variants, we were able to determine which sections of the tertiary structure of the INSR protein were linked to type 2 Diabetes Mellitus. Ultimately, we created a map of the INSR protein and indicated which parts of the protein structure had significant effects on type 2 Diabetes Mellitus. We concluded differences in the charged/polar amino acids on the lower right section of the structure(amino acid numbers 487 and 750) were associated with pathogenic variants, while charged/polar differences on the lower left of the structure(amino acid numbers 669 and 784) had no impact on pathogenicity. We observed several differences in the amino acid chain structure, however they were not unique to the pathogenic variants; they were also seen in the benign variants.

## Introduction

Proteins conduct a wide variety of cellular functions such as: cell signalling, cell division, metabolism, communication, and transport(1). Although DNA encodes how to build these proteins, proteins conduct many of the important functions in humans(1). The structure of a protein depends on their physical structure and chemical composition, such as polarity, charge, and the ability to form bonds(1). The function of proteins can be determined through comparison to amino acid chains of known proteins(2).

Although DNA codes the protein, a variety of folding, bonding and cleaving processes occur(3). Originally, scientists believed the interactions involved in protein folding were described in the amino acid sequence(3). Recent research, however, has shown the interaction of chaperone proteins has an effect on protein folding(3). Hence, an important part of bioinformatics research is not just studying the DNA sequences but also the protein sequences and their protein models.

The INSR protein is involved in the signal transduction pathway for insulin. The INSR gene produces a receptor tyrosine kinase(4). When insulin binds to this receptor, a signal transduction pathway commences, and glucose enters the cell(4). Certain diseases associated with insulin insensitivity such as type A insulin resistance syndrome are related to the INSR gene(4). Further, type 2 diabetes, the type of diabetes associated with insulin resistance, have been researched as a contributor to type 2 diabetes(5). As such, in this article we will investigate the structure of the INSR protein.

We collected FASTA protein sequences of pathogenic variants(subjects with type 2 diabetes) and benign variants(subjects without type 2 diabetes). We used Swiss-Model, a protein homology modeling tool, to compare the models against a “base” model, which was the FASTA sequence obtained from insulin receptor isoform X2. We then compared the structures generated from Swiss-Model. We cataloged the differences between the benign variants and the base and the pathogenic variants and the base.

## Materials and Methods

We obtained the FASTA protein sequences from the database SNP residing in NCBI. We named the variants as they were named when submitted to SNP. We collected seven pathogenic variants and six benign variants. From these variants, we submitted them to Swiss-Model under their names as they appeared in SNP. Through using the structure comparison tool in the Swiss-Model, we compared the variants and the base. We took several pictures of the differences in structure and cataloged the difference in FASTA sequences.

## Results

We compared the structure of the INSR protein in the base protein and benign proteins, and the base protein and pathogenic protein. We defined two types of differences: structural differences and amino acid property differences. Structural differences are defined as differences in the structure of the protein, while amino acid property differences are differences in the amino acids, causing the corresponding section of the protein to have different properties.

In the seven benign protein variants we observed, we saw the following number of differences with the base model amino acid abbreviation and location number on the left and benign model amino acid abbreviation and location number on the right.

**Table 1.**

| Base model | Benign model |
| --- | --- |
| ASP 261 | GLU 261 |
| GLN 303 | HIS 303 |
| ASP 546 | GLU 546 |
| PHE 669 | LEU 669 |

Some of these amino acid variations caused amino acid property differences, with the base model amino acid abbreviation, location, and changing property on the left and the benign model amino acid abbreviation, location, and changing property on the right:

**Table 2.**

| Base model | Benign model |
| --- | --- |
| GLN 303 not aromatic | HIS 303 aromatic |
| PHE 669, not aliphatic | LEU 669, aliphatic |
| PHE 669, aromatic | LEU 669, aliphatic |
| GLN 699, neutral | GLU 699, negatively charged |

On the INSR model, the location of these differences are seen in the figures below:

**Figure 1.**
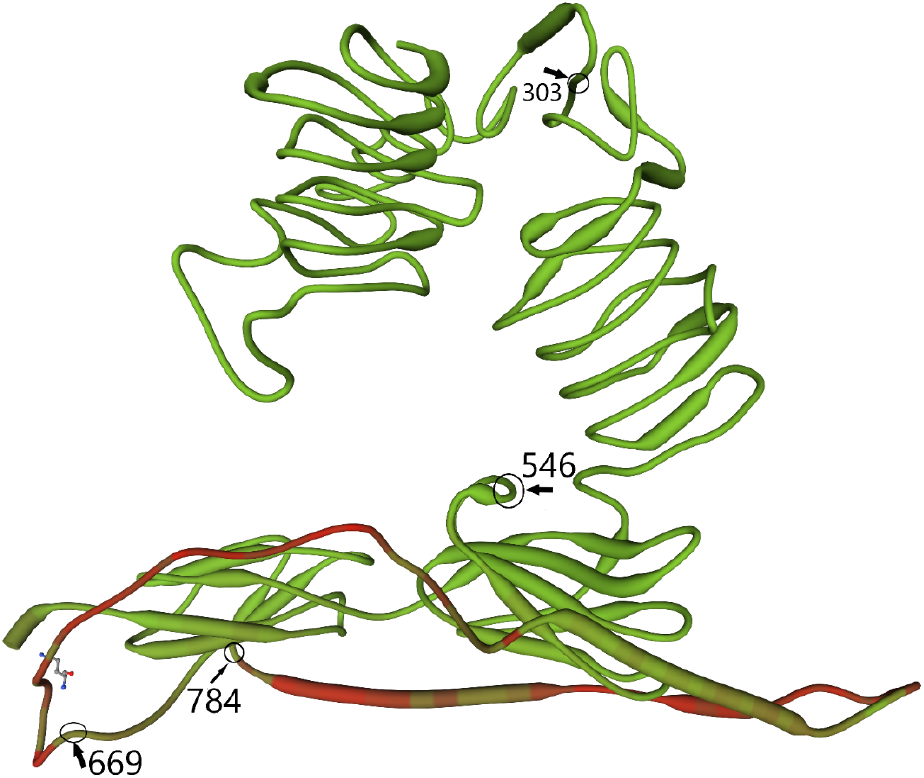

**Figure 2.**
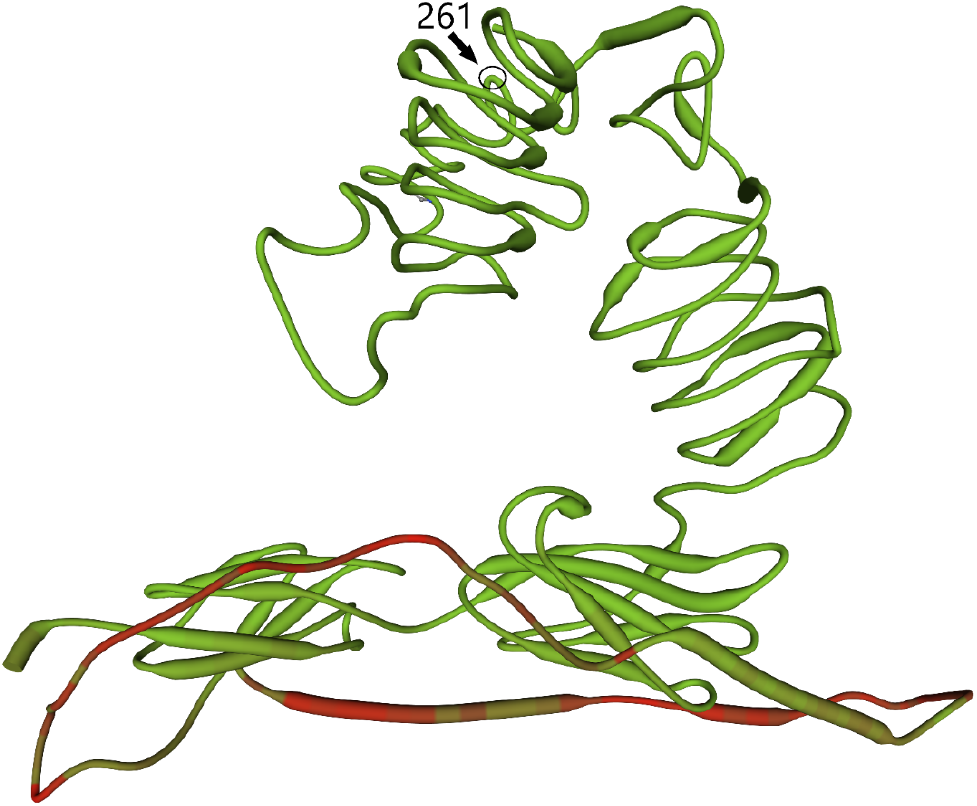

We also observed structural differences in the benign variants:

We can see from figure 3 the amino acid chain for the base and benign variants split between amino acid 544 and 556, 668 and 719, and 743 and 779. In the split from 668 to 719, we see the base chain loops outward to the left and loops over the benign variant. On the other hand, the variant chain turns to the right quicker and loops one time. In the 743 to 799 split, the benign variant shoots up, then falls down while the base variant moves in a relatively straight line. In the 544 to 556 split, the benign variant moves to the right, while the base moves to the left.

**Figure 3.**
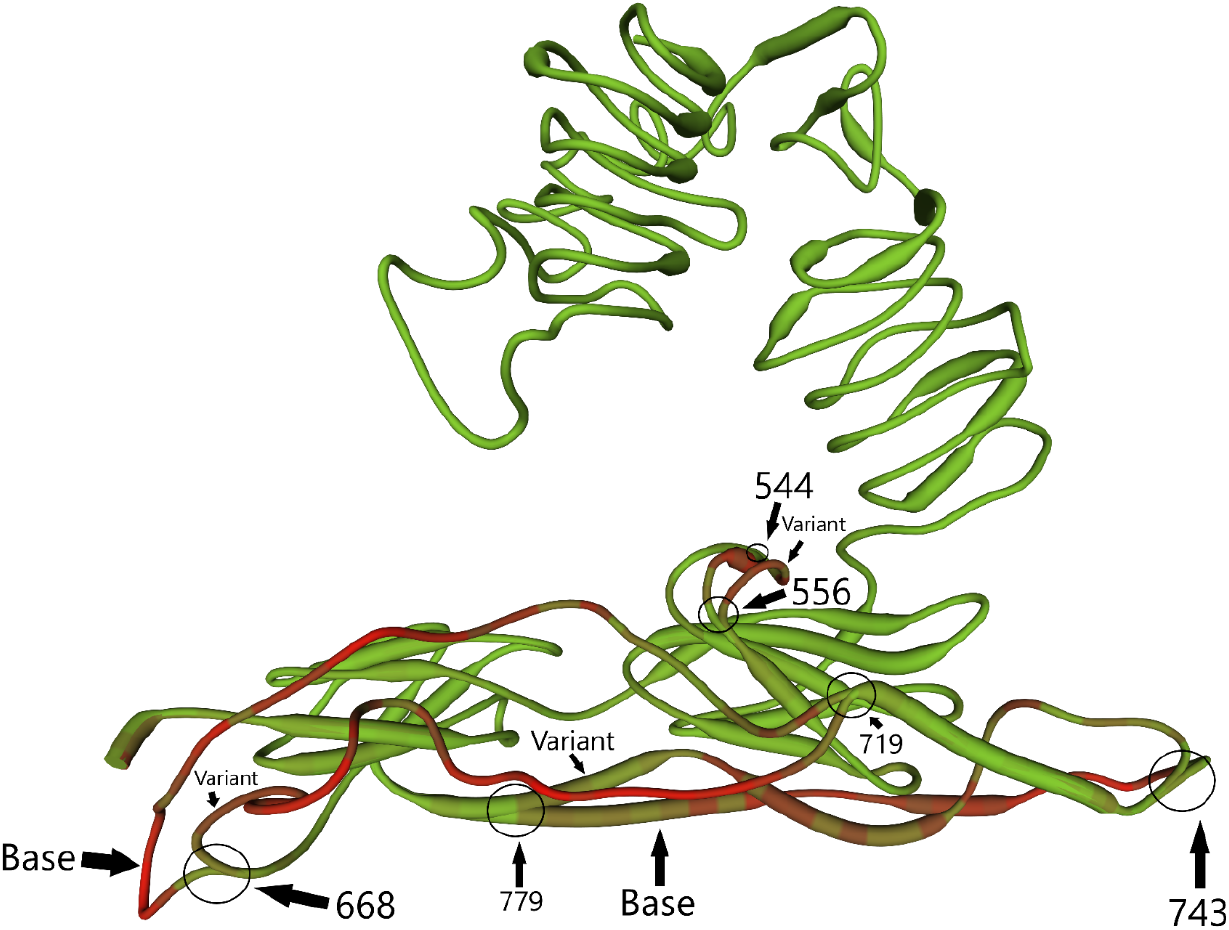

In the pathogenic variants, we see the following amino acid differences with the amino acid and location on the base on the left and the corresponding difference in the pathogenic variant on the right:

**Table 3.**
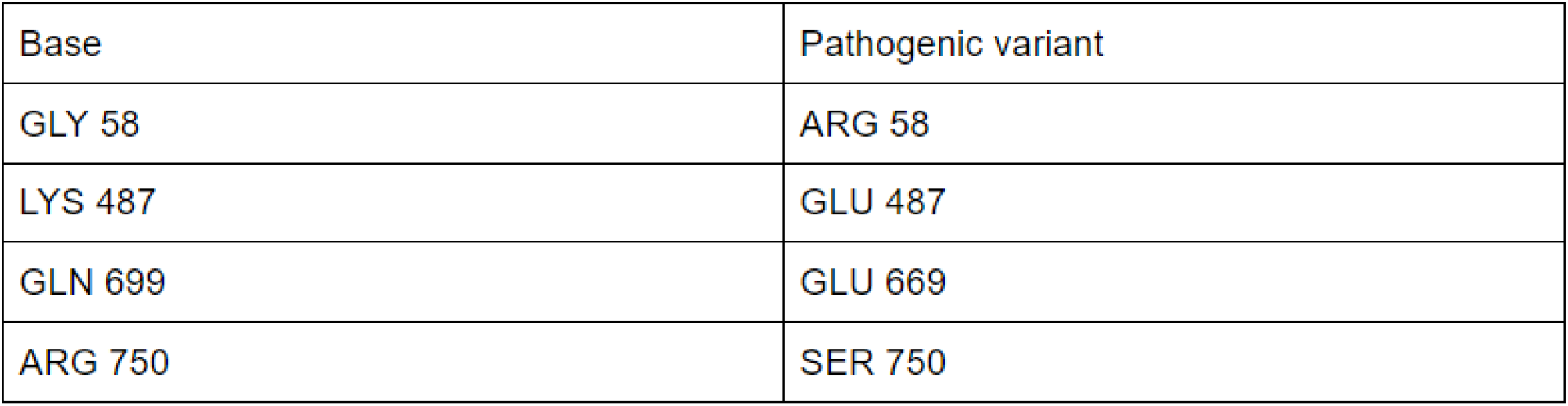

| Base | Pathogenic variant |
| --- | --- |
| GLY 58 | ARG 58 |
| LYS 487 | GLU 487 |
| GLN 699 | GLU 669 |
| ARG 750 | SER 750 |

The amino acid property differences we observed in the pathogenic differences due to these amino acid differences are:

**Table 4.**

| Base | Pathogenic variant |
| --- | --- |
| GLY 58, non polar | ARG 58, polar |
| GLY 58, neutral | ARG 58, positive charge |
| LYS 487, positive charged | GLU 487, negative charged |
| GLN 699, neutral | GLU 699, negative charge |
| ARG 750, positive charge | SER 750, neutral |

The location of these differences on the INSR protein structure are seen in the figure below:

**Figure 4.**
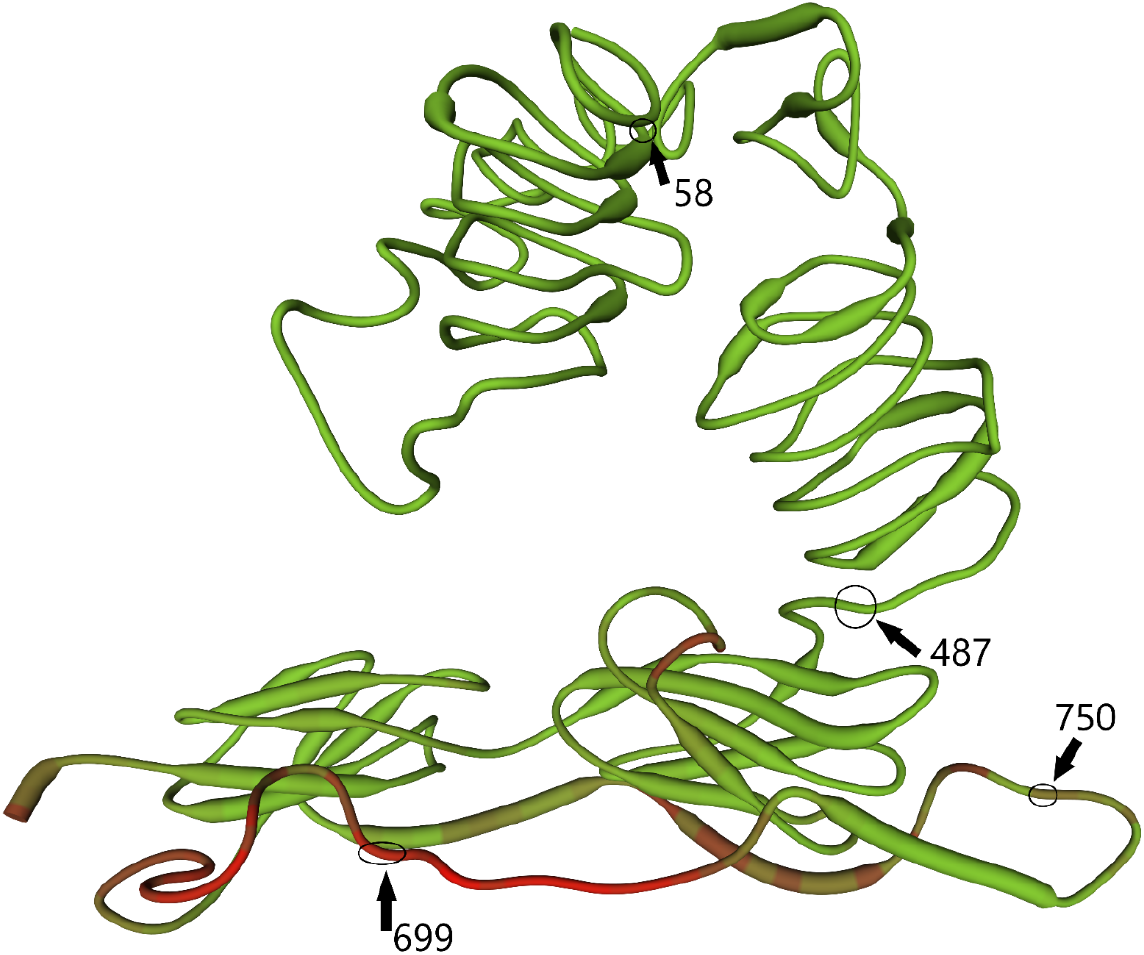

We observed similar structural differences in the pathogenic variants:

## Discussion

We compared the differences in INSR protein structure between benign and pathogenic variants. We predict the development of insulin resistance is related to differences in the structure of the INSR protein. Differences we observed in the benign variants don’t have an effect, while differences observed in the pathogenic variants do.

From the benign versions, we see the ability to form aliphatic bonds in location 669, aromatic bond in location 303, charged/polar amino acids in location 699, and neutral amino acids in location 784 are different from the base model.

We see the presence of charged/polar amino acids in location 58 and 699, while the presence of neutral amino acids in location 750 and a flipped charge in location 487 in the pathogenic variations. In the figure below, we show the location of all the differences observed, with the benign and pathogenic variations.

**Figure 5.**
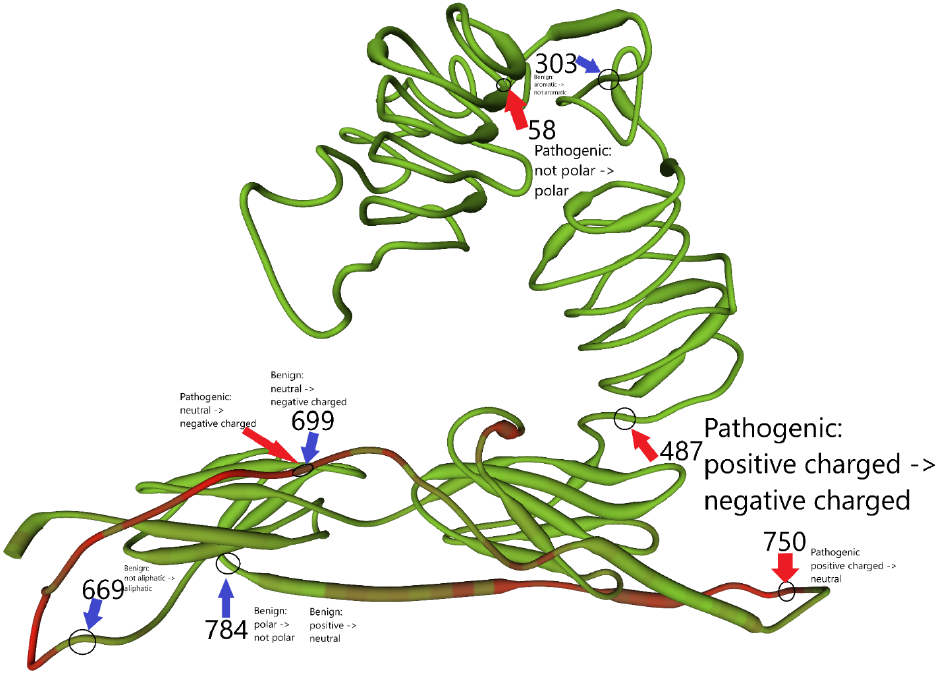

From this model we see alterations in the left bottom section of the protein are mostly benign. The change in aliphatic nature of amino acid 669 seems to have no effect, and the changes in charge/polarity in amino acid 784 also have no effect. From this, we can hypothesize the lower left section of the INSR protein is not involved in binding to insulin, as changes in 669’s and 784’s bonding nature and polarity/charge would have an effect on the binding of insulin. The change on 699 is more interesting. A change from neutral to negative is found in both pathogenic and benign variants. Therefore, we can conclude that it is not involved in the pathogenesis type 2 diabetes, but rather a change which has a null effect.

The alterations on the lower right of the protein are both pathogenic. These alterations transform amino acid 487 from positive to charged and amino acid 750 from positive to neutral. Changing the charge of both these amino acids are associated with pathogenic variants, so we predict the lower right section of the protein plays a greater role in the binding to insulin.

In the upper middle section of the protein, we see two variations, one pathogenic and one benign. The pathogenic variant transforms from polar to non polar. As the amino acid seems to be buried between other structures, we predict this nonpolar amino acid is sheltered on the inside of the protein structure, away from water. In the pathogenic form, we predict this amino acid destroys the structure of the protein as it is polar and hence attracting water. The other variant is the bening change of 303 from aromatic to not aromatic. As this change is benign, we predict it does not affect insulin binding, and hence it seems insulin does not bind to the top part of the protein structure.

We observed similar structural differences between the benign and pathogenic variants. In both the benign and pathogenic variants, we saw splits in the amino acid chains from 544 to 556, 668 to 719 and 743 to 779. Hence, we suppose these differences were not associated with pathogenic results, as they are also present in benign variants.

## Bibliography

1. Breda, Ardala. “Protein Structure, Modelling and Applications.” Bioinformatics in Tropical Disease Research: A Practical and Case-Study Approach [Internet]., U.S. National Library of Medicine, 14 Sept. 2007, www.ncbi.nlm.nih.gov/books/NBK6824/.

2. Alberts, Bruce. “Analyzing Protein Structure and Function.” Molecular Biology of the Cell. 4th Edition., U.S. National Library of Medicine, 1 Jan. 1970, www.ncbi.nlm.nih.gov/books/NBK26820/.

3. Cooper Geoffrey M. “Protein Folding and Processing.” The Cell: A Molecular Approach. 2nd Edition., U.S. National Library of Medicine, 1 Jan. 1970, www.ncbi.nlm.nih.gov/books/NBK9843/.

4. “INSR Insulin RECEPTOR [Homo Sapiens (Human)] - Gene - NCBI.” National Center for Biotechnology Information, U.S. National Library of Medicine, www.ncbi.nlm.nih.gov/gene/3643.

5. Bodhini D; Sandhiya M; Ghosh S; Majumder PP; Rao MR; Mohan V; Radha V; “Association of His1085his INSR Gene Polymorphism with Type 2 Diabetes in South Indians.” Diabetes Technology & Therapeutics, U.S. National Library of Medicine, pubmed.ncbi.nlm.nih.gov/22775283/.

